# Abi3 regulates microglial ramification and dynamic tissue surveillance *in vivo*

**DOI:** 10.1101/2021.03.19.436147

**Authors:** Elena Simonazzi, Ruth E. Jones, Fangli Chen, Adam Ranson, Joshua Stevenson-Hoare, Valentina Escott-Price, Frank Sengpiel, B. Paul Morgan, Philip R. Taylor

## Abstract

A rare coding variant of Abelson-interactor gene family member 3 (Abi3) is associated with increased risk of late-onset Alzheimer’s Disease (AD). Although Abi3 is recognised as a core microglial gene, its role in microglia is largely unknown. Here we demonstrate that Abi3 is crucial for normal microglial morphology, distribution, and homeostatic tissue surveillance activity *in vivo*.

## Main

The actin cytoskeleton is essential for diverse macrophage cellular processes – including polarisation, adhesion and migration – and its rapid remodelling is particularly important in microglia, where processes constantly form, retract and re-form to monitor the brain parenchyma at a velocity calculated at 1.47 µm/min^1^. This highly dynamic actin cytoskeleton is essential for microglia to perform their role in maintaining brain homeostasis by phagocytising debris and dead cells, pruning synapses and reacting to changes in the brain microenvironment following tissue damage, presence of pathogens or protein aggregates such as the β-Amyloid plaques in Alzheimer’s Disease (AD)^2^. To achieve this plasticity, the actin cytoskeleton is finely controlled by small GTPases, such as Rac^3^, and their downstream effectors, including the WASP-family verprolin homologous (WAVE) proteins^4,5^. Of particular interest is the WAVE2 complex, which plays a key role in the regulation of lamellipodia formation in fibroblast and macrophage migration^6,7^.

This pentameric complex consists of WAVE2, Nap-1, HSPC300, and PIR121/Sra-1^8^ and members of the Abelson-interactor family^9,10^. While ABI1 is the most common and well-studied member, in the brain ABI3 is predominantly expressed by microglia^11^ and considered a signature microglia gene^12,13^. ABI1 and ABI3 are known to be mutually exclusive within the WAVE2 complex^14^ and exogenous ABI3 expression reduced WAVE2 translocation to the cell periphery^15,16^ by preventing c-ABL/WAVE2 phosphorylation^15^. Ectopic expression of ABI3 in NIH3T3 fibroblasts, which endogenously express extremely low levels of ABI3^14^, has been shown to impair cell spreading^14^. In a cancer cell line ABI3 forced overexpression caused cell cycle arrest in G0/G1 phase^17^, while in cultured rat hippocampal neurons both overexpression and knock-down of Abi3 caused changes in dendritic spine morphogenesis and synapse formation^18^. Moreover, ABI3 was found to be enriched in microglial clusters around Aβ-plaques in AD brains^19^ and recent whole-exome microarray analyses identified a rare ABI3 coding variant, p.Ser209Phe, as a risk factor for AD development^20^, prompting a need to understand its role in microglia, a cell type which has been implicated in AD^21^. Taken together, these observations suggest a role for ABI3 in microglia in AD, which has not yet been explored. Given the requirement for a physiologically-relevant microenvironment for microglial phenotype, this study focused on the study of cells of the macrophage lineage and microglia *in vivo*, both with endogenous Abi3 expression levels.

Macrophage motility, as well as their phagocytic activity, depend on the ability of these cells to correctly form lamellipodia and filopodia structures. Interestingly, ABI3 overexpression in NIH3T3 fibroblasts altered the formation of lamellipodial and actin-mediated cell spreading^14^. To examine Abi3 role in a more physiologically-relevant context, we generated conditionally-immortalised macrophage precursor (MØP) cell lines by transduction of CD117^+^ bone marrow cells from age and gender matched Abi3-WT and -KO mice with a retrovirus encoding oestrogen-dependent version of Hoxb8^22^. Abi3 deficiency was confirmed at genomic level in both Abi3-KO mice and MØP cell lines and by qPCR of macrophages derived from the MØPs (Supplementary Fig. 1a-b). Macrophages differentiated from MØPs in M-CSF resemble bone-marrow derived macrophages^22^ and exhibited increased Abi3 expression in Abi3-WT cells, but expression was not evident in the Abi3-KO cells, as expected (Supplementary Fig. 1c-d). The MØP-derived macrophages enabled examination of cell spreading in an appropriate cell type as well as in the context of physiologically-relevant endogenous Abi3 expression levels. The ability of Abi3-WT and -KO macrophages to spread on fibronectin-coated coverslips was tested using an actin-mediated spreading assay. Cells were examined at four time points over a 4-hour period and analysed (Supplementary Fig. 2) to obtain area and solidity values for the spreading of each cell. Abi3-deficient macrophages showed a marked increase in cell size compared to Abi3-WT control cells, with a noticeable reduction in the number of ramifications (Fig. 1a). Indeed, Abi3-WT cells had significantly smaller surface and were richer in processes, with a resulting decrease in cellular solidity, at all time-points (Fig. 1b-i and Supplementary Table1), even up to a week after re-plating (Supplementary Fig. 3). Similar results were obtained with two additional pairs of independently generated cell lines, confirming that the observed phenotype was linked to genotype and not an artifact of an individual cell line.

**Fig. 1.**
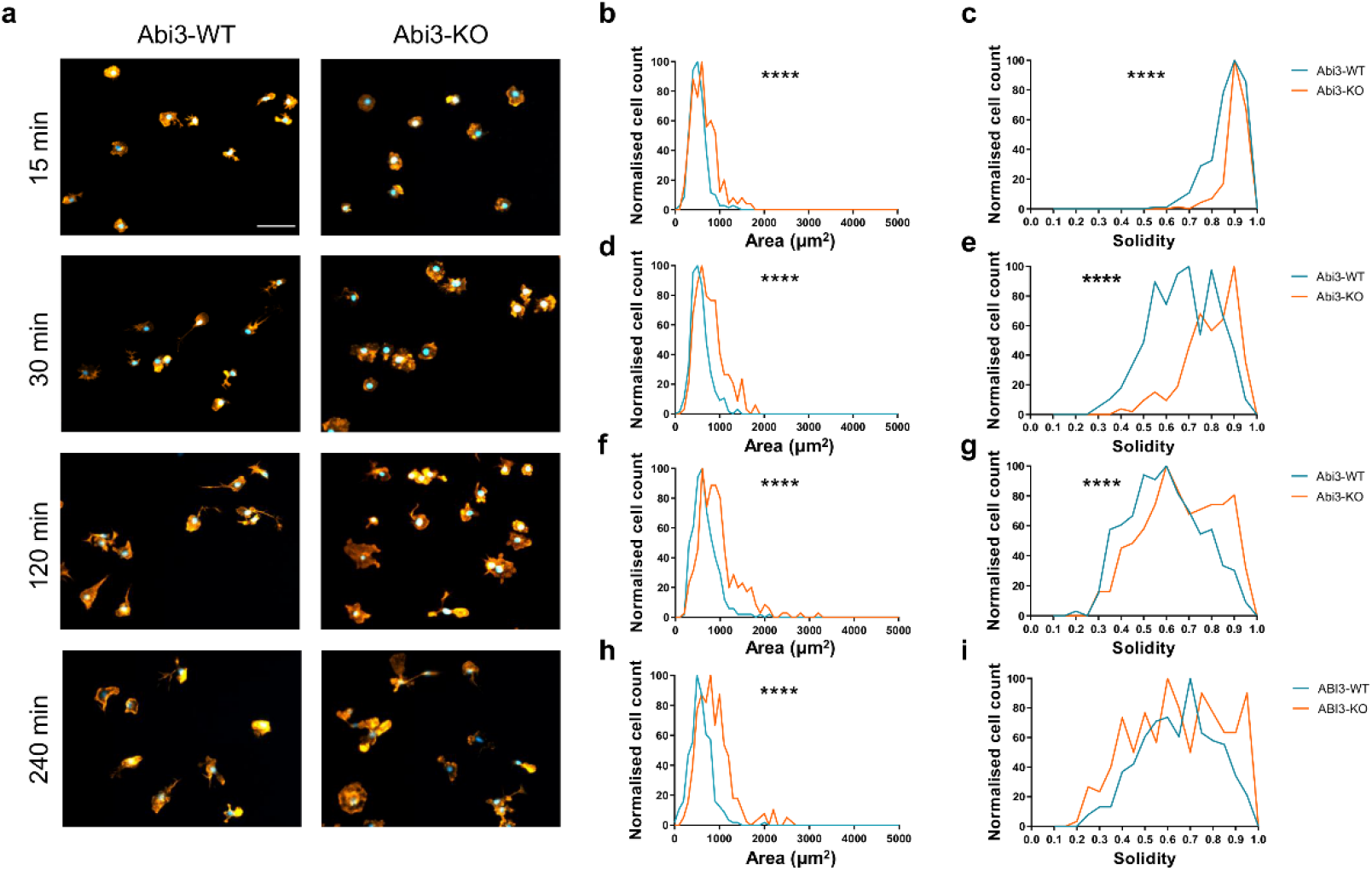
Alterations in Abi3 levels significantly impact the actin-mediated spreading of macrophages. **a**, Representative images of actin cytoskeleton staining (Alexa Flour 555-labelled Phalloidin) on M-CSF differentiated macrophages derived from Abi3-WT or KO conditionally-immortalized precursor cell lines at the indicated time points (15, 30, 120 and 240 minutes) during an actin-mediated spreading assay; scale bar 100 μm. **b-i**, Representative quantification of area (**b, d, f** and **h**) and solidity (**c, e, g** and **i**) values at each time point of the spreading assay on Abi3-WT and -KO macrophages, shown as normalized histograms of the frequency distribution of the samples; blue line indicates cells derived from the Abi3-WT mice, while the orange line those from the Abi3-KO mice. Data were analysed via Mann-Whitney test; ****p≤0.0001; n>140 (see Supplementary Table 1). This experiment was repeated three times, with a total of three independent cell lines (see Supplementary Table 1), with similar results.

Given the marked phenotypic differences observed between Abi3-WT and -KO macrophages and the high expression of Abi3 in microglia, we hypothesized that Abi3 could similarly control microglial morphology and associated functionality. As microglia are intrinsically connected to astrocytes and neurons, any perturbation of this equilibrium could have additional repercussions, especially in a pathological brain microenvironment, as observed in AD; for this reason, we examined the microglia of the animals *in situ*.

Microglial morphology and distribution were assessed in the Prefrontal Cortex (PFC) and the Hippocampus of young (8-week-old) Abi3-WT and –KO mice due to the contribution of these two brain regions to memory formation and retention^23,24^ and their involvement in AD pathogenesis^25,26^. In accordance with the *in vitro* macrophage data, Abi3-KO microglia stained with an Iba1 antibody lacked the complex ramifications of Abi3-WT cells, and consistently showed shorter and thicker processes in both brain regions (Fig. 2a). Abi3-KO mice also showed a significant increase in microglia number and concomitant reduced Nearest Neighbour Distance (NND) in the PFC (Fig. 2b-c) as well as in the Hippocampus (Fig. 2e-f) when compared to the Abi3-WT mice. However, the Iba1^+^ area of brain tissue was decreased compared to the Abi3-WT controls in both PFC (Fig. 2d) and Hippocampus (Fig 2g) despite the increased cell density, suggesting an impairment in microglia ability to efficiently monitor brain parenchyma.

**Fig. 2:**
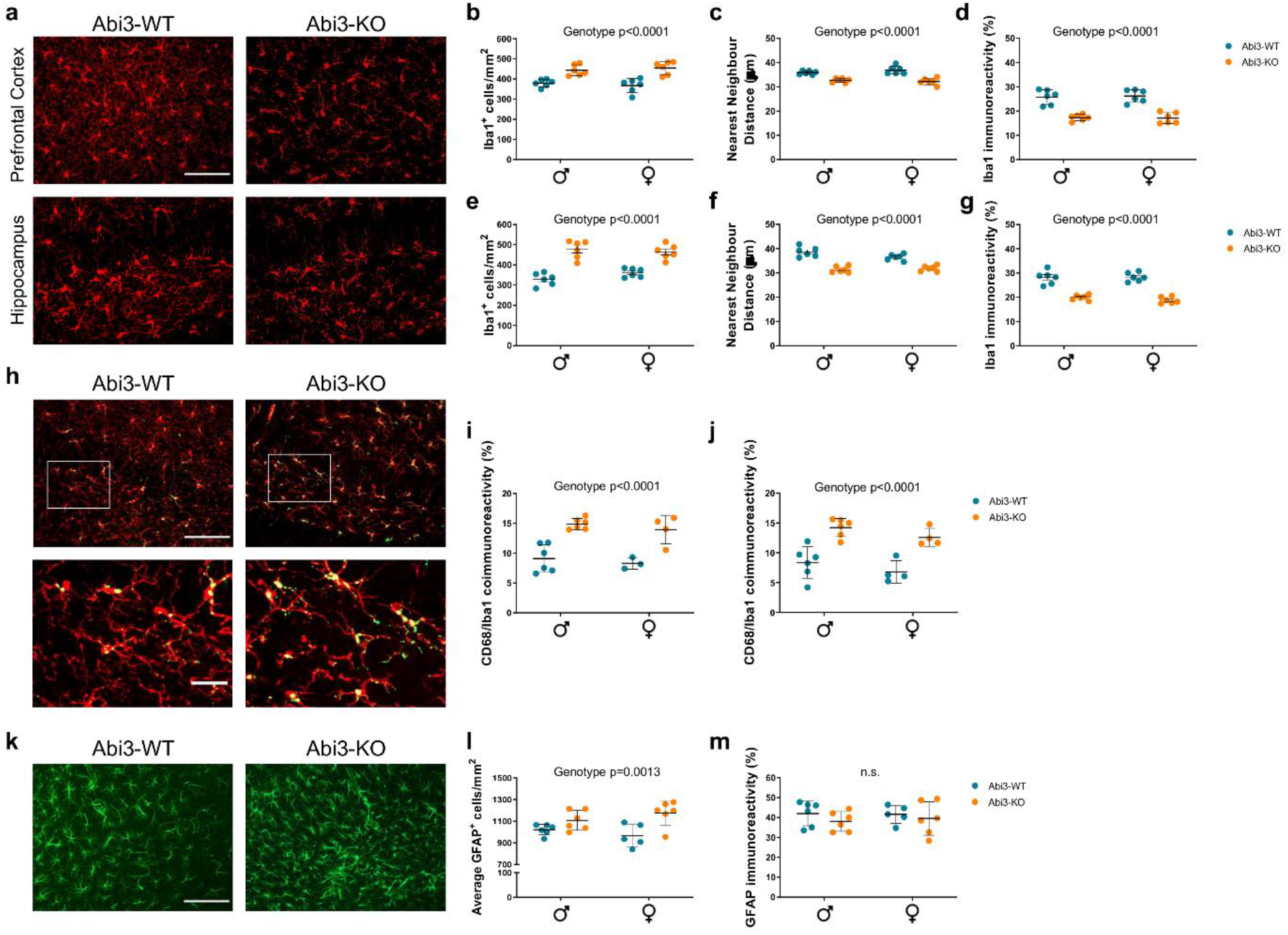
The absence of Abi3 leads to significant alterations of glia in the brain. **a**, Representative pictures of the Iba1 staining performed on the Prefrontal Cortex (PFC) and the Hippocampus of Abi3-WT and -KO mice, showing a marked reduction of immunoreactivity in Abi3-KO mice. Scale bar 100 μm. **b-g**, Quantification of **b/e**, microglia density, **c/f**, Nearest Neighbour Distance (NND) and **d/g**, Iba1 immunoreactivity in Abi3-WT and -KO mice PFC and Hippocampus (**b-d** and **e-g**, respectively). n=6/group. **h**, Representative pictures of the Iba1 (red) and CD68 (green) co-staining performed on the Hippocampus of Abi3-WT and -KO mice. Scale bars 100 μm (top row) and 25 μm (bottom row). **i-j**, Quantification of CD68 and Iba1 colocalization in the **i**, PFC and **j**, Hippocampus of Abi3-WT and -KO mice n=6 for males; n=3-4 for females. **k**, Representative images of Abi3-WT and -KO astrocytes stained with an anti-GFAP antibody in the Hippocampus. Scale bar 100 μm. **l-m**, Quantification of **l**, astrocytes density and **m**, GFAP immunoreactivity withing the Hippocampus of Abi3-Wt and -KO mice. n=6/group, except Abi3-WT females (n=5). In all the graphs, blue symbols indicate Abi3-WT mice, orange ones Abi3-KO animals. Each circle represents the average value of one animal; the black horizontal lines represent the mean of each experimental group ± SD. Data were analysed via Two-way ANOVA test for all the experiments; the p-value of the Genotype effect identified by the Two-way ANOVA is reported on each graph.

The gross phenotype observed in Abi3-KO mice has been reported as indicative of activated microglia. CD68 is a lysosomal marker upregulated in phagocytic microglia and regarded as a marker of microglia activation^27^. For this reason, we investigated CD68 levels in the absence of Abi3. Interestingly, histological analysis of Abi3-WT and -KO mice brains highlighted a significant increase in CD68 levels within microglia in the absence of Abi3 in both the hippocampus (Fig. 2h, j) and PFC (Fig. 2i).

Activated microglia have been shown to release multiple factors – including pro-inflammatory cytokines, Nitric Oxide, Transforming Growth Factor α and Vascular Endothelial Growth Factor B – which in turn regulate astrocytic phenotype^28^. Thus, while Abi3 is mainly expressed by microglia^11^, the loss of Abi3 could indirectly impact astrocytes. We therefore assessed Glial Fibrillary Acidic Protein (GFAP) positive cells within the Hippocampus of young (8-week-old) Abi3-KO mice. We observed an increase in the number of GFAP^+^ astrocytes within the hippocampus of Abi3-KO mice (Fig. 2k-l), while no significant difference was evident in the levels of GFAP immunoreactivity (Fig. 2m), suggesting a potential alteration of astrocytic morphology following loss of Abi3.

To examine in detail the extent of the morphological abnormalities presented by Abi3-KO microglia, we traced the 3-dimensional structure of single microglia (Fig.3a) and analysed the resultant data using a linear mixed model (Supplementary Table 2 and 3). Compared to the Abi3-WT, this analysis revealed a significant increase in branch segments length (Fig. 3b), due to reduction in branch ramification, and an increase in ramification diameter (Fig. 3c), consequently affecting the branch volume (Fig. 3d) in the Abi3-KO mice. Abi3-KO cells also presented a significant decrease in total number of branching points (Fig. 3e) as well as in number of consecutive branching points (Fig. 3f) compared to the Abi3-WT mice. The difference in ramifications between the two genotypes of mice was highlighted by the Sholl profiles of the four experimental groups, with both male and female Abi3-KO mice showing a dramatic decrease in Sholl intersections (Fig. 3g-h). Overall, these differences lead to a significantly smaller cell surface (Fig. 3i) and volume (Fig. 3j) in the absence of Abi3.

**Fig. 3:**
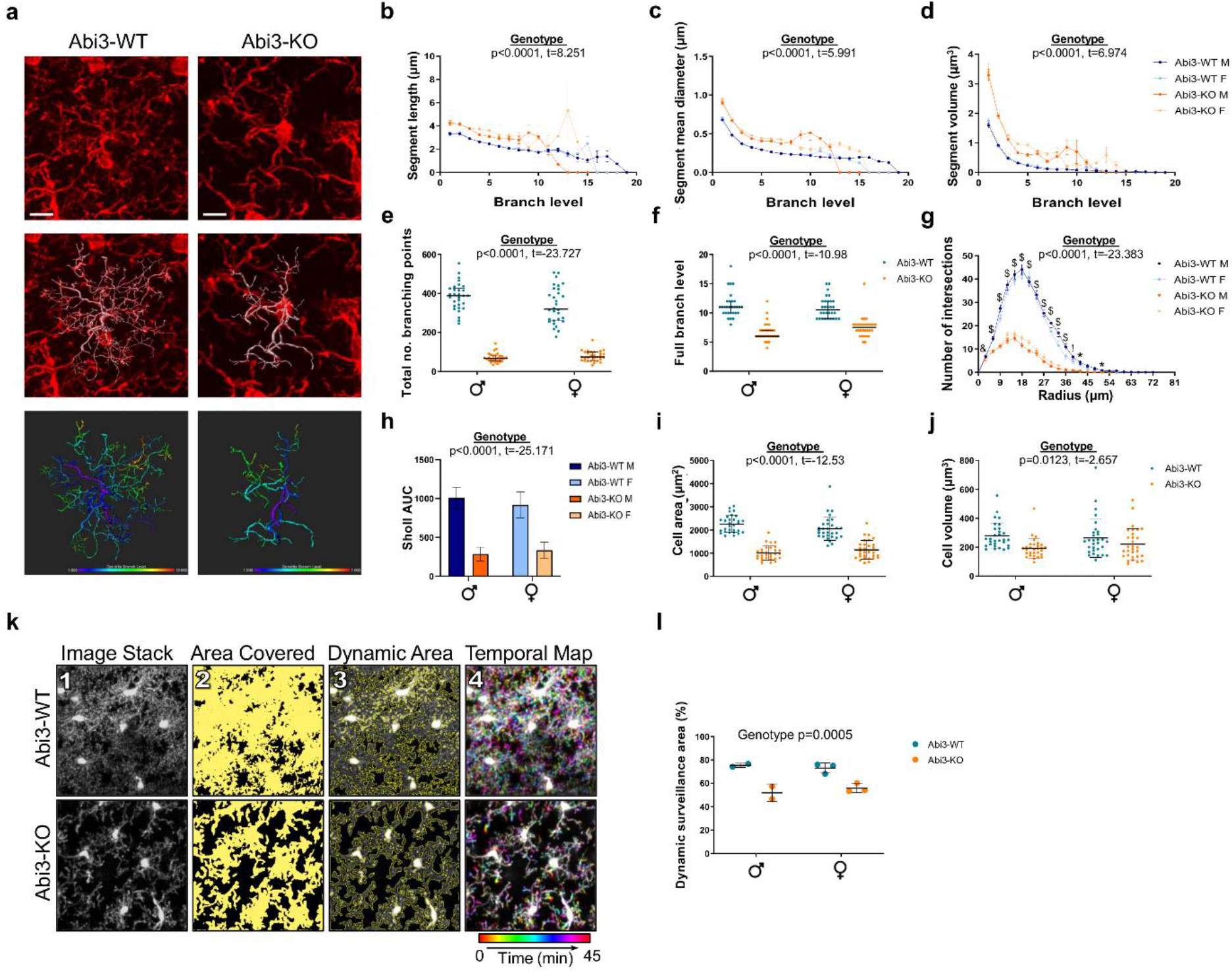
Abi3 knock-out severely impacts microglia functionality in vivo due to its detrimental effects on the ramification apparatus of these cells. **a**, Representative images of 3D-traced microglial ramifications of single cells in the CA1 region of the Hippocampus of Abi3-WT and -KO mice using Imaris. Scale bar 10 μm. Each individual cell, stained with an anti-Iba1 antibody (top row, red) was manually 3D traced (middle row, white) on Imaris; bottom row presents an example of the same tracing colour-coded based on branch level to highlight the difference in arborisation complexity between genotypes. **b-d**, Quantification of **b**, ramification length, **c**, mean diameter, and **d**, volume of segments forming Abi3-WT and –KO microglial ramifications. n>9000 segments from 30 cells/group. **e-h**, Evaluation of: **e**, the total number of branching points; **f**, overall branching level; **g**, Sholl profile and **h**, consequent Area Under the Curve of male and female animals for each genotype; **i**, cell surface; **j**, cell volume. n=30 cells/group. Data were analysed using a linear mixed-model including genotype and sex – and wherever appropriate, branch level or Sholl radius – as fixed effects and animal IDs as random effect; t- and p-values for the Genotype effect are reported on each graph and in Supplementary Table 2 and 3. For the Sholl analysis, the individual p-values relative to the Genotype effect are reported on the graph by means of symbols: * p≤0.05, & p≤0.01, ! p≤0.001, $ p≤0.0001**k**, Schematic representation of the analysis workflow on orthogonal projections originated from a 45-minute-long time stack acquired on awake Abi3-WT and –KO mice by two-photon microscopy. After generating an orthogonal projection of the time stack (1), a threshold was applied to select the total area covered by microglia soma and processes (2) and the ‘static’ (and therefore brighter) regions were subtracted, producing the dynamic area ROI (3) outlined in yellow on top of the orthogonal projection of the time stack to demonstrate the accuracy of the analysis; the temporal map (4) allows visual appreciation of the rapid actin remodelling and tissue coverage in Abi3-WT microglia during the 45-minute acquisition period. **l**, Quantification of the percentage of area dynamically surveyed by microglia ramifications in Abi3-WT and –KO mice brain over a period of 45 minutes. Data were analysed via Two-way ANOVA and p-value relative to the Genotype effect is reported on the graph. In all the graphs, Abi3-WT mice are indicated by blue symbols, Abi3-KO by orange ones; in the histograms, dark colours represent male animals, while lighter colours indicate females. For **b, c, d**, and **g**, each datapoint represents the average obtained from all the cells belonging to the same experimental group, ± S.E.M.; for **h**, each bar represents the average of all the cells in the group ±SD; in the remaining graphs, each datapoint represents the value relative to a single cell and the black horizontal lines represent the median of each experimental group ± interquartile range (**e** and **f**) or the mean ± SD (**i, j**, and **l**).

These histological findings pointed to a likely alteration of microglia surveillance activity, due to the marked decreased in ramification complexity in the absence of Abi3. Microglial surface protrusions are extremely dynamic and are constantly scanning the surrounding parenchyma, and this can be visualised *in vivo* by 2-photon microscopy through cranial windows in awake mice^29^. Abi3-WT and –KO mice were crossed with a Cx3cr1-EGFP reporter line^30^, to generate mice heterozygous for the EGFP reporter. Two-photon analysis on unchallenged awake mice revealed that Abi3-KO microglia exhibited reduced surveillance of brain parenchyma, with only 51.88% ± 7.49% and 53.32% ± 3.77% (for males and females respectively; mean ± SD) of the total area covered by motile microglial processes during the 45-minute acquisition period, compared to 75.41% ± 1.80% and 73.07% ± 4.35% in Abi3-WT animals (Fig. 3k-l and Supplementary Video 1 and Video 2).

To date, the physiological function of Abi3 in macrophages, and specifically microglia, was largely unknown. Previous *in vitro* studies implicated it in the regulation of actin dynamics^14–16,18,31^. This is the first report to address the role of Abi3 in microglia *in vivo*, demonstrating its control of both morphology and function of these cells.

The dystrophic phenotype observed in the Abi3-KO mice, together with their increased CD68 levels, is of particular interest in the pathological context of AD, where microglia have been implicated in Aβ clearance through phagocytosis^21^, and an impairment in microglia migration towards amyloid plaques has been shown to significantly worsen amyloid burden^21^. Moreover, alterations in microglial phenotype could affect their synaptic pruning activity, possibly leading to the increased synapse loss observed in AD^32^. These observations highlight the need for further investigation of the role of microglial motility and tissue surveillance in the risk of AD.

In conclusion, the data presented here support the hypothesis that Abi3, a core microglial gene, contributes to AD development through its fundamental role in the regulation of microglial morphology and movement, which alters homeostatic surveillance of brain tissue by microglia. This could in turn cause an impairment in microglial phagocytic activity – with potentially detrimental effects on debris removal and Aβ clearance – as well as synaptic pruning. Therefore, our studies identify Abi3 as a key molecule of interest for further investigation in AD.

## Acknowledgments

E.S. is funded by an UK Medical Research Council (MRC) GW4 BioMed Doctoral Training Partnership studentship. P.R.T is funded by a MRC UK Dementia Research Institute Professorship and a Wellcome Trust Investigator Award (107964/Z/15/Z). BPM is funded by a MRC UK Dementia Research Institute Professorship. We would like the thank the staff of our animal facilities for the care of the animals used in this study. For the purpose of Open Access, the author has applied a CC BY public copyright licence to any Author Accepted Manuscript version arising from this submission. The datasets generated during and/or analysed during the current study are available from the corresponding author on reasonable request. Unique reagents will be made available to reasonable scientific request, where permitted by existing intellectual property constraints.

## Author information

### Author contributions

E.S. and R.E.J. performed experiments and analysed data. F.C. performed craniotomies and acquired two-photon videos. A.R. wrote the MATLAB code employed in two-photon experiments, while J.S.-H. provided the code for Imaris data analysis. E.S, R.E.J., V.E.-P., F.S, B.P.M. and P.R.T. provided intellectual contributions. E.S, R.E.J and P.R.T wrote the paper. All authors reviewed the manuscript.

## Methods

### Animals

Male and female transgenic Abi3-KO mice (Abi3^tm1.1(KOMP)Vlcg^; Jax) and wild type control mice (Abi3-WT) were used in these experiments. Abi3-KO mice arrived as heterozygous and were intercrossed to generate lines of WT and KO mice. Abi3-KO mice were co-housed with Abi3-WT animals of the same sex from weaning. For histological studies, animals were sacrificed at 8-9 weeks of age. For two-photon experiments, Abi3-WT and –KO mice were crossed to B6.129P2(Cg)-Cx3cr1^tm1Litt^/J mice^30^ (Jax) and only GFP-heterozygous mice were used for the following analysis. All animal experiments were approved by the Animal Welfare and Ethical Review Body (AWERB) - sub group of the Biological Standards Committee - and conducted in accordance with UK Home Office Guidelines and Animal [Scientific Procedures] Act 1986 which encompasses EU Directive 2010/63/EU on the protection of animals used for scientific purposes. Animals were housed in conventional open top cages with environmental enrichment under controlled conditions (20–24°C, 12-hour light/dark cycle, food and water *ad libitum*).

Genotyping of Abi3-WT and -KO mice was performed using standard PCR conditions (30 amplification cycles at 59 °C) and the following primers (all 5’→3’): WT-Fw ACCCAGATCCCTGAGAATTTG; WT-Rv CAAGTCCTGAAGGGAGAACG; KO-Fw CGGTCGCTACCATTACCAGT; KO-Rv CAGCCCAAGAGGTAGACAGG. The final PCR products were 334 base pairs (bp) in Abi3-WT mice and 473 bp in Abi3-KO mice. Cx3cr1 genotyping was performed by touchdown PCR (10 cycles with declining annealing temperature from 65 °C to 60 °C, followed by 20 cycles at 60 °C) using the following primers (all 5’→3’): WT-Fw GTCTTACGTTCGGTCTGT; Mt-Fw CTCCCCTGAACTGAAAC; Common-Rv CCCAGACACTCGTTGTCCTT. The final PCR products were 410 bp for WT mice and 500 bp for Cx3cr1-GFP mutant mice.

### Generation of conditionally-immortalized Macrophage Precursor (MØP) cell lines

*Hoxb8* conditionally-immortalised macrophage precursor (MØP) cell lines were derived from CD117^+^ enriched bone-marrow cells taken from individual 8week-old mice, euthanised with CO_2_ asphyxiation, using established methods^22^. Six MØP lines were generated in total, an initial paired Abi3 WT and KO lines derived from 8 week-old female mice followed by four lines, one for each genotype and gender generated from individual 8 week-old mice. Three pairs of polyclonal lines were generated to ensure that observed changes were reproducible and not an artifact of a specific cell line. In the absence of β-estradiol and the presence of 20 ng/mL Macrophage Colony-Stimulating Factor (M-CSF; Peprotech) MØP cells differentiate into cells phenotypically similar to bone marrow-derived macrophages^22^.

### Cell culture

All cells were maintained at 37°C in a humid incubator with 95% air 5% CO_2_. Abi3-WT and -KO MØPs were maintained in 6-well plates with Roswell Park Memorial Institute 1640 (RPMI; Gibco®, ThermoFisher) supplemented with 10% heat-inactivated Foetal Bovine Serum (FBS, Gibco®, South America) and 100 U/ml Penicillin, 10 μg/ml Streptomycin (Gibco®, ThermoFisher), 1 μM oestrogen (stock: 10 mM β-estradiol in absolute ethanol; Sigma) and 10 ng/mL recombinant murine Granulocyte-Macrophage Cytokine Stimulating Factor (GM-CSF; Peprotech).

### Genomic KO PCR

Genomic DNA (gDNA) was isolated from 2 × 10^6^ MØP cells in accordance with the instructions of the GenElute™ Mammalian Genomic DNA Miniprep Kit (Sigma). Separate PCR reactions were prepared using the GoTaq® Master Mix (Promega) and primers targeting Abi3 exon 5-7 (5’→3’ Fw: GATCCCTGAGCCGGTG; Rv: GGGCAACTCTGCTT) or exon 8 (5’→3’ Fw: TCTTCAGCACCTGTTGC; Rv: AGCACGATTAACTGAGAGA) per manufacturer’s specifications. In brief, the following PCR conditions were used:

- Abi3 exon 5-7 was amplified by touchdown PCR (10 cycles with declining annealing temperature from 55°C to 45°C, followed by a further 20 cycles at 45°C).
- Abi3 exon 8 was amplified by touchdown PCR (10 cycles with declining annealing temperature from 55°C to 50°C, followed by a further 20 cycles at 50°C).

The final PCR products were resolved on a 2% Agarose gel at 100V for 30 minutes. The predicted band sizes were 766 bp for exon 5-7 PCR and 962 bp for exon 8.

### qPCR

Approximately 2 × 10^6^ undifferentiated MØP cells or M-CSF differentiated MØP cells were lysed in accordance with the Qiagen RNeasy Mini kit (Qiagen). The concentration of RNA was measured using either a Nanodrop™ 2000 (Thermofisher Scientific) or a DS-11 FX+ spectrophotometer (DeNovix®). RNA was then converted to complementary DNA (cDNA) using the Precision™ Reverse-Transcription Premix 2 kit (PrimerDesign LTD) per manufacturer’s directions. qPCR reactions were performed using the 2X Precision®FAST qPCR Master Mix (with SYBR-green and low ROX) according to manufacturer’s instructions (PrimerDesign LTD). The following primers were used (5’→3’): *Abi3* Fw: TCAAAACCCAGCAGGCTCCC; Rv: CTTGTCTGTGGCCTGCAAGTAGT; *Ywhaz* Fw: TTGAGCAGAAGACGGAAGGT; Rv: GAAGCATTGGGGATCAAGAA. Plates were then run on either a Viia™ 7 or QuantStudio™ 7 Flex Real-Time PCR Systems (both Applied Biosystems™) with 40 repetition of data collection steps at 60 °C. *Abi3* mRNA expression data was assessed using the x40 cycle and ΔΔCT method where *Ywhaz* was used as an endogenous control gene using the QuantStudio™ Real Time PCR software. Three biological samples were taken from each undifferentiated MOP cell line and their M-CSF differentiated counterpart (n=3 per group).

### Spreading assay

Sterile 13 mm coverslips were coated with 50 µg/ml fibronectin (Sigma) in Hank’s Balanced Salt Solution (HBSS; Gibco®, ThermoFisher) for a minimum of 40 minutes at room temperature. Triplicate coverslips were prepared for each experimental group. M-CSF differentiated Macrophages were lifted by incubation for 10 minutes at 37°C with Accumax (Sigma). Cells were counted using Muse® Cell Analyzer (Merk Millipore) and plated onto 24-well plates containing the FN-coated coverslips and 500 µl of complete fresh RPMI media (supplemented with 20 ng/mL M-CSF) in each well. For the two short timepoints, 15 and 30 minutes, 10^4^ cells were plated in each well while for the longer incubations (2 and 4 hours) 5 × 10^3^ cells were used to prevent overcrowding. At each time point the media was discarded and cells were washed once with DPBS, before being fixed with 4 % Paraformaldehyde (PFA; Sigma) for 10 minutes at room temperature. Cells were then washed with DPBS, permeabilised for 3 minutes at room temperature with 0.1 % Triton-X-100 (Sigma) in DPBS, washed again as before and blocked with 1 % Bovine Serum Albumin (BSA; Sigma) in DPBS for 20 minutes at room temperature. F-actin was stained for 20 minutes at room temperature with fluorescent Phalloidin (Invitrogen, ThermoFisher) conjugated to Alexa Fluor (AF) −555 or −647 at a 1:40 dilution in 1% BSA in DPBS. Finally, cells were washed, incubated 5 minutes with at room temperature with 500 ng/ml 4’,6-Diamidino-2-Phenylindole Dilactate (DAPI; ThermoFisher) in DPBS and mounted in ProLong®Gold (ThermoScientific).

Coverslips were imaged using a Zeiss Apotome Axio Observer microscope (Zeiss) or an EVOS™ FL Auto 2 Imaging System (Thermo Scientific) microscope with a 20x objective. Slides were independently blinded and 15-30 random fields were analysed for each experimental group (10 for each coverslip) using Fiji software (Fiji is just ImageJ, version 1.52p^33^). After a first step of brightness and contrast optimisation, manual separation of cells in close contact was performed using a 2 μm-thick black line. After carefully adjusting the threshold and generating a binary file, the “Fill holes” tool automatically filled any gap within the particles, which were then analysed using the “Analyse Particles” function. Due to the expected ramified morphology of macrophages, the two chosen parameters for the following statistical analysis were cell area and cell solidity (described by the equation: [Area]/[Convex Area]). The experiment was repeated a total of 3 times: twice on the original pair of cell lines, and one more time on two additional pairs in order to exclude cell line-specific artefacts.

### Brain extraction and preparation

8 weeks old mice were euthanised with intraperitoneal overdose of Euthatal® (Merial Animal Health Ltd) followed by intracardial perfusion-fixation with 20 ml ice cold DPBS and 50 ml ice cold 4 % PFA. Brains were removed from the skull and left over-night in 4 % PFA at 4 °C. Brains were then transferred and stored in DPBS with 0.1 % Sodium Azide (Sigma) at 4 °C until further processing. Samples were then cut into 50 µm-thick free-floating coronal sections using a Leica VT1200S vibratome (Leica Biosystems) and stored long-term at 4 °C in DPBS with 0.1 % Sodium Azide.

### Immunofluorescent staining

Coronal sections from two regions of interest, the Prefrontal Cortex (between 3.08 mm to 2.58 mm from the bregma) and the Hippocampus (−1.34 mm to −2.18 mm from the bregma), were selected for each mouse using Images from the Mouse Brain atlas^34^as a visual reference. Sections were washed with 1x Tris-buffered saline (TBS; 137 mM NaCl, 20mM Tris, pH 7.6) and then permeabilized and blocked in TBS supplemented with 0.5 % Triton X-100, 1 % BSA and 0.3M Glycine (FisherScientific) for 1 hour at room temperature. Samples were then incubated over night at 4°C with anti-Iba1 (AF635-conjugated rabbit monoclonal, 1:100; Wako), anti-GFAP (chicken monoclonal, 1:1000; Abcam) and/or anti CD68 (rat monoclonal, 1:100; Bio-Rad) antibodies in 0.5 % Triton X-100, 1 % BSA in TBS. Sections were washed 3 times with 1x TBS and incubated for 2 hours at room temperature with appropriate secondary antibodies (goat anti-chicken FITC, 1:200, Abcam or goat anti-rat AF488, 1:200, The Jackson Laboratories) in 0.5 % Triton X-100, 1% BSA in TBS. After 3 more washes, the brain slices were counterstained for 10 minutes at room temperature with 500 ng/ml DAPI in DPBS. After a last wash, the samples were mounted in ProLong®Gold and stored in the dark at 4 °C.

### Image acquisition and analysis

Brain sections were imaged using a Zeiss Cell Observer Spinning Disk confocal microscope (Zeiss) with a 20x objective. Eight z-stacks were acquired for the PFC and 14 for the Hippocampus region. The central 30 optical sections of each z-stacks were transformed into a frontal maximum-intensity orthogonal projection and used for the following analyses with Fiji Software (Fiji is just ImageJ, version 1.52p^33^). After utilising the “Background subtraction” tool, Iba1 or GFAP positive cells were manually counted using the “Cell Counter” plugin. Care was taken not to include cells with their cell body partially outside the edges of the image. Each image was then converted to the 8-bit format and Iba1 or GFAP positive pixels were selected by applying a threshold to generate a binary image. The “Measure” tool allowed the evaluation of area percentage covered by Iba1^+^ or GFAP^+^ pixels. In the case of the microglial staining, the 8-bit image was further processed to obtain Nearest Neighbour Distance (NND) values. Two sequential filters from the MorphoLibJ plugin library were applied to the images: a Gray Scale Attribute opening filter (Operation = “Opening”, Attribute = “Area”, Minimum Value = 25 pixels, Connectivity = 8) was applied to isolate the cells from the background, followed by an opening Morphological filter (Operation = “Opening”, Element = “Octagon”, Radius = 1 pixel) in order to separate the cell body from the ramifications. After generating binary images using the “Entropy Threshold” plugin, missing cell bodies or wrongly included ramifications were manually corrected to reduce bias. Centroids were calculated for each cell body selecting a 30µm minimum size filter in the “Analyse Particles” tool. Finally, the “NND” plugin was used to calculate the relative values for all the identified cell bodies. Experiments were performed in blinded conditions, in order to avoid any operator bias.

For CD68 and Iba1 co-localisation analysis, the area percentage covered by Iba1^+^ cells was used to generate a ROI, that was then applied to the CD68 staining (processed in the same manner as the Iba1 channel to obtain a binary image) before using the “Measure” tool, in order to make sure only colocalizing pixels were comprised in the analysis.

### Morphometric analysis of single cells

Iba1^+^ cells in the CA1 region of the Hippocampus were images using a Zeiss Cell Observer Spinning Disk confocal microscope (Zeiss) with a 40x oil immersion objective. Between 4 and 6 z-stacks of random fields were acquired for each mouse. Single cells were carefully cropped out of each image taking care to select only cells that were entirely comprised within the z-stack and whose cell body was not overlapping with adjacent cells. A total of 5 cells per animal was then analysed using the Filament Tracer module of Imaris software (Bitplane, Oxford Instruments, version 9.3.1). The filament starting point was placed at the soma and the seed point threshold was adjusted to include all the ramifications. Manual editing of the algorithm-generated filament was performed by an operator blinded to the experimental condition, in order to remove or correct ill-traced ramifications and to add any branch missed by the algorithm. The numerical output of the following parameters was extracted for further statistical analyses: branch length, branch mean diameter, branch volume, total number of branching points, full branch level, cell area, cell volume and Sholl intersections.

### Two-photon microscopy

A week before the imaging session, 8-9 weeks old Abi3-WT and –KO mice crossed to B6.129P2(Cg)-Cx3cr1^tm1Litt^/J mice underwent a craniotomy as described by Goldey et al.^29^. Briefly, a 3 mm-large circular portion of skull centred over the primary visual cortex (3.00 mm lateral to lambda and 1.64 mm anterior to the lambdoid suture^29^) was removed and replaced with two 3 mm cover glasses and one 5mm cover glass on the top, and a metal head plate fixed to the skull by dental cement. A week after the surgery mice were image, d using a resonant scanning two-photon microscope (Thorlabs, B-Scope) with a 16x 0.8 NA objective with 3-mm working distance (Nikon). Mice were awake during the imaging session and positioned on a cylindrical treadmill. Laser power was set to 30mW (at the sample) for all recordings. Each mouse underwent a single imaging session, during which a total of 30 z-stacks (comprised of 25 optical sections, divided by a 2µm step size for a total z volume of 50 μm) of the same random area were acquired every 90 seconds over a period of 45 minutes using a 7x zoom in order to ensure visualisation of the thin processes. Each focal plane of the z-stack was imaged 10 consecutive times before moving to the next one, which then allowed image registration to be performed with a custom written software in MATLAB^®^ (MathWorks). Each z-stack was then merged in a maximum intensity orthogonal projection and then collated with all the others, in chronological order.

The resulting file was then used to perform a quantification of the area covered by Abi3-WT and –KO microglia using Fiji software. A single orthogonal projection was obtained flattening the time-stack. A first threshold was applied to select all the ramifications, while a second one was used to only comprise the brightest objects, which mostly encompassed the cell bodies. The percentage of “dynamic area” – i.e. the area covered only by dynamic structures – was measured after subtracting the second binary image from the first one.

### Statistics

Statistical analyses were mostly performed using GraphPad Prism (GraphPad Software, version 8). The M-CSF-differentiated Macrophage spreading assay was analysed by Mann-Whitney test. All n and p values are indicated in the relevant figure legend and Supplementary Table. Two-way ANOVA tests for genotype and sex factors were used for basic microglia, CD68 and astrocyte characterisation, as well as for the data generated from the analysis of the two-photon videos. Data generated from Imaris were analysed using linear mixed-model analysis in R (version 4.0.0). To account for possible confounders, in addition to the genotype the regression models included sex and mice IDs, as well as radius or branch level wherever appropriate; genotype, sex and radius/branch level were included as fixed effects, while the animal’s ID was included as a random effect to account for the animal-specific variability.

