## Supplementary for "Abi3 regulates microglial ramification and dynamic tissue surveillance *in vivo*"

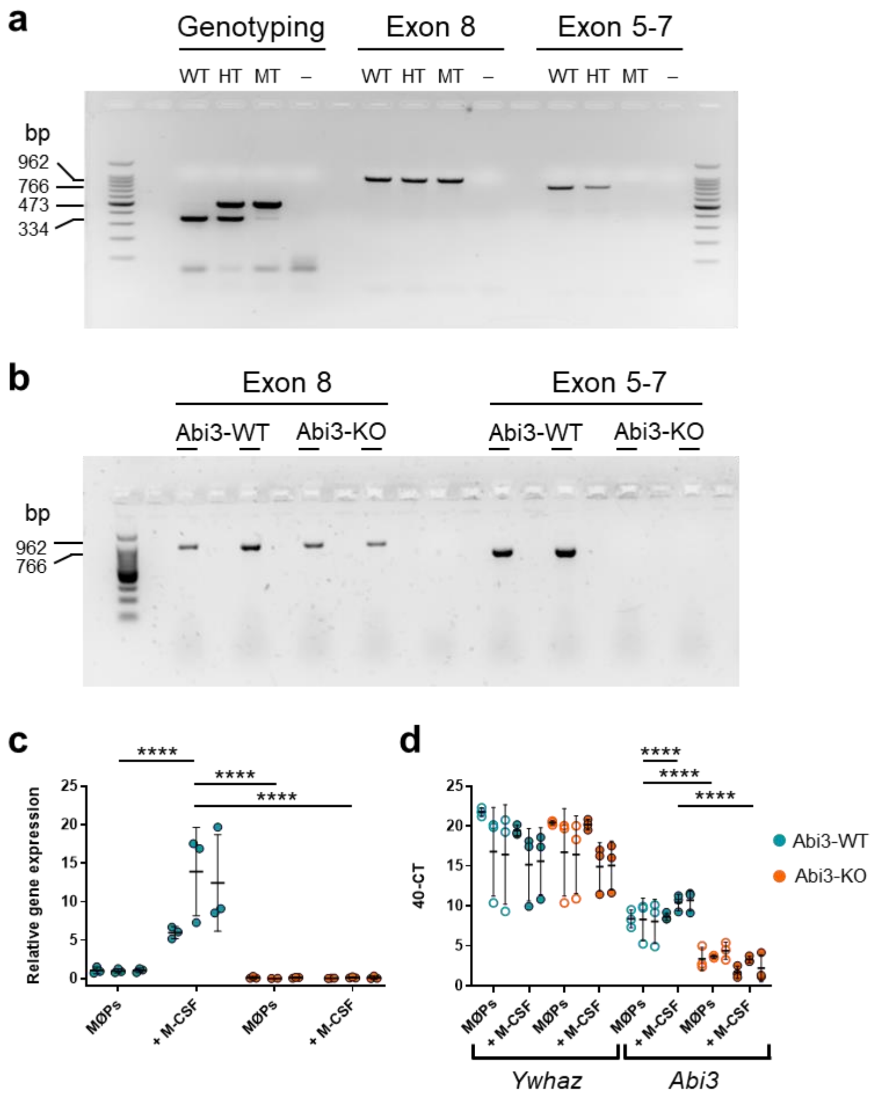


***Supplementary Fig. 1: Abi3 knock-out confirmation in Abi3-KO mice and MØP cell lines. a-b,*** *Representative genomic PCR of Abi3 Exon 5-7 and Exon 8 performed on DNA extracted from* ***a,*** *Abi3-WT and -KO mice and* ***b,*** *MØPs confirming the absence of Exon 5-7 in Abi3-KO animals and cells.* ***c,*** *Abi3 mRNA relative expression as analysed by qPCR using YWHAZ mRNA levels as endogenous housekeeping control. Cells deriving from 3 different cell lines for each genotype were tested after being either maintained with GM-CSF and β-Estradiol (MØPs) or differentiated using M-CSF. Data were normalised to the average value of the WT MØPs. The relative gene expression confirmed the absence of Abi3 mRNA in Abi3-KO cells.* ***d,*** *Graphical representation of the 40-CT values deriving from the qPCR analysis. Blue dots represent Abi3-WT cells, while orange indicates Abi3-KO samples. Each dot represents the result of n=3 separate experiments. The horizontal black lines indicate the mean ± SD. Data analysed by One-Way ANOVA; ****$p \leq 0.0001$.*

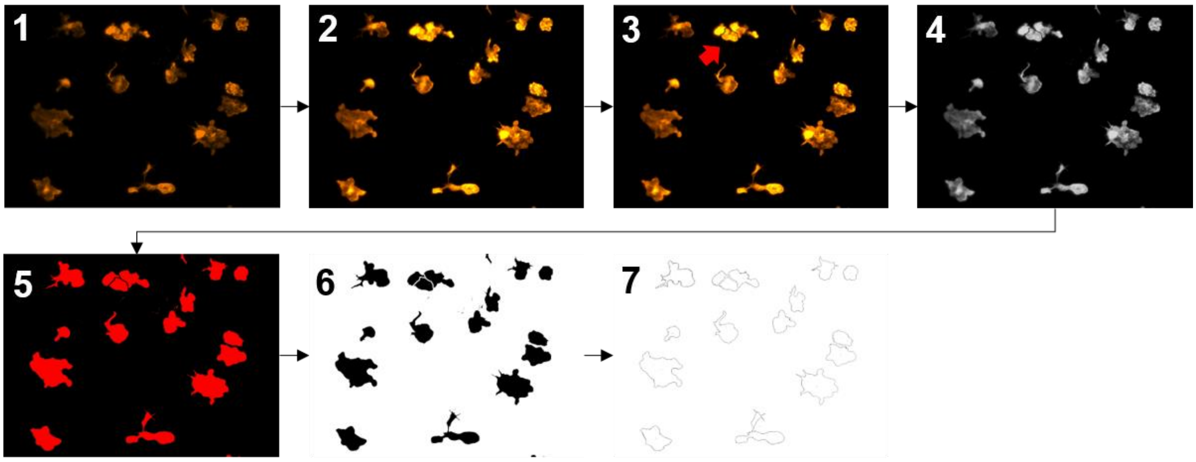


***Supplementary Fig. 2: Schematic representation of the analysis workflow of M-CSF differentiated macrophages spreading assay.*** *The original TIFF images of the Phalloidin staining (1) were imported into Fiji software and processed by adjusting brightness and contrast (2). Adjacent cells were then manually separated with a 2 µm-thick black line (3, red arrow) and the images were converted to the 8-bit format (4). Finally, after a threshold was applied (5) and any artificial gap in the deriving binary images was filled using the "Fill holes" function (6), cells were analysed using the "Analyse particles" tool, excluding particles present on the borders or with a surface of less than 20 µm*$^{2}$ *(to avoid including debris or background pixels), generated outlines images of the analysed cells (7).*

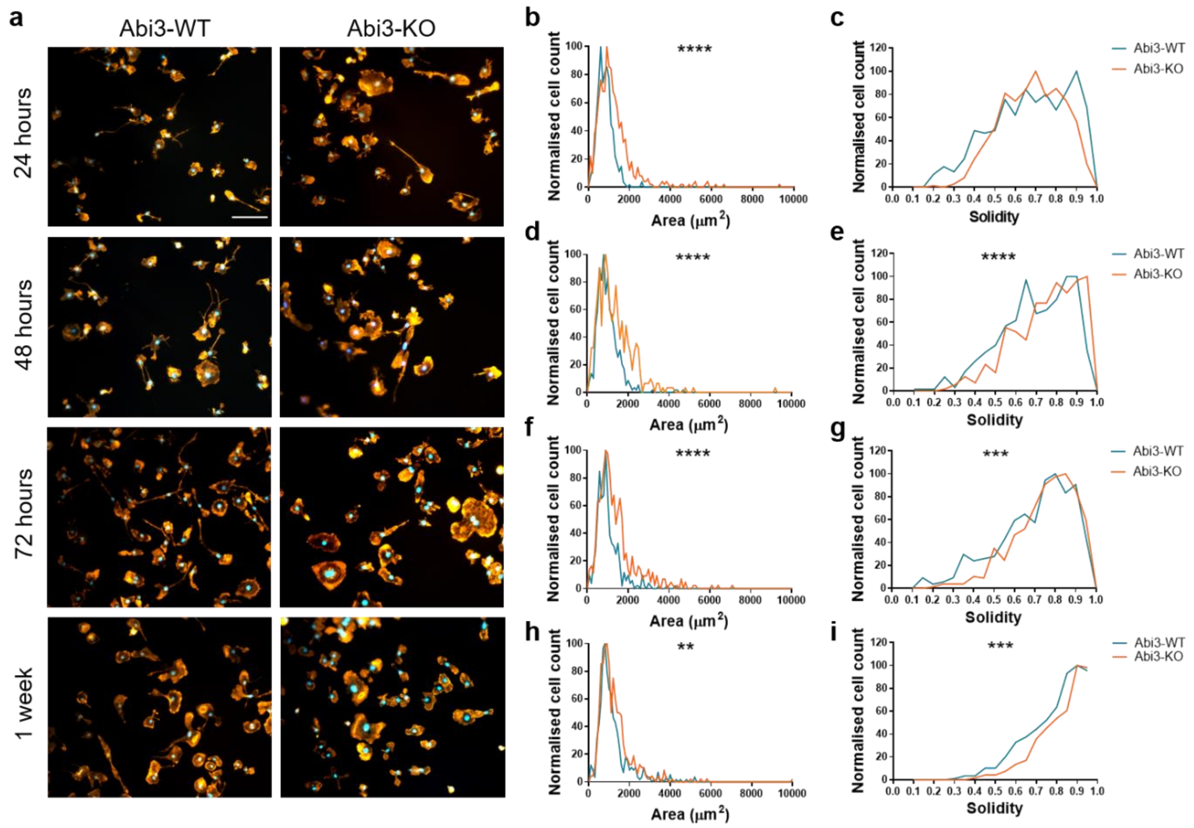


***Supplementary Fig. 3: The effects of Abi3 absence on macrophage spreading ability last for up to a week after replating. a,*** *Representative images of M-CSF differentiated Abi3-WT and -KO macrophages, stained with AF555-Phalloidin, after 24, 48, 72 hours and 1 week adhesion; scale bar 100 μm.* ***b-i,*** *Representative quantification of area (**b, d, f** and **h**) and solidity (**c, e, g** and **i**) values at each time point of the spreading assay, shown as normalized histograms of the frequency distribution of the samples. Abi3-WT cells are indicated in blue, Abi3-KO in orange. Data were analysed via Mann-Whitney test; ** $p \leq 0.01$, *** $p \leq 0.001$, **** $p \leq 0.0001$; n>350.*

**Area — 15 min**

| Cell line pair | A | | A | | B | | C | |
|---|---|---|---|---|---|---|---|---|
| Genotype | Abi3-WT | Abi3-KO | Abi3-WT | Abi3-KO | Abi3-WT | Abi3-KO | Abi3-WT | Abi3-KO |
| Number of values | 221 | 287 | 286 | 140 | 613 | 517 | 449 | 552 |
| Minimum | 86.05 | 86.57 | 81.46 | 160.5 | 22.89 | 39.63 | 49.25 | 86.57 |
| 25% Percentile | 317.2 | 265.9 | 402.2 | 436.7 | 485.9 | 613.7 | 460.3 | 456.2 |
| Median | 398.3 | 357.6 | 501.1 | 605.2 | 582.7 | 762.2 | 562.3 | 551.0 |
| 75% Percentile | 511.9 | 531.3 | 609.2 | 814.9 | 703.1 | 949.4 | 659.6 | 672.2 |
| Maximum | 1402 | 1738 | 1426 | 1734 | 1946 | 2552 | 1501 | 1869 |
| Mann-Whitney test | n.s. | | **** | | **** | | n.s. | |

**Area — 30 min**

| Cell line pair | A | | A | | B | | C | |
|---|---|---|---|---|---|---|---|---|
| Genotype | Abi3-WT | Abi3-KO | Abi3-WT | Abi3-KO | Abi3-WT | Abi3-KO | Abi3-WT | Abi3-KO |
| Number of values | 385 | 402 | 291 | 227 | 595 | 538 | 454 | 429 |
| Minimum | 77.81 | 104.1 | 58.27 | 244.8 | 21.35 | 22.32 | 37.90 | 206.2 |
| 25% Percentile | 572.2 | 528.9 | 420.6 | 544.8 | 474.4 | 596.2 | 445.4 | 495.3 |
| Median | 736.9 | 748.7 | 534.3 | 712.6 | 601.6 | 767.7 | 557.1 | 597.9 |
| 75% Percentile | 960.8 | 1062 | 655.1 | 945.7 | 759.5 | 991.9 | 684.9 | 737.0 |
| Maximum | 2621 | 2706 | 1448 | 1837 | 2674 | 3004 | 1557 | 3851 |
| Mann-Whitney test | n.s. | | **** | | **** | | **** | |

**Area — 120 min**

| Cell line pair | A | | A | | B | | C | |
|---|---|---|---|---|---|---|---|---|
| Genotype | Abi3-WT | Abi3-KO | Abi3-WT | Abi3-KO | Abi3-WT | Abi3-KO | Abi3-WT | Abi3-KO |
| Number of values | 432 | 422 | 272 | 261 | 392 | 392 | 324 | 359 |
| Minimum | 82.45 | 82.45 | 245.4 | 240.8 | 22.32 | 21.35 | 20.39 | 141.8 |
| 25% Percentile | 484.2 | 565.8 | 477.2 | 621.9 | 513.8 | 619.0 | 507.7 | 489.6 |
| Median | 680.7 | 856.7 | 606.0 | 864.4 | 665.5 | 872.5 | 655.7 | 660.6 |
| 75% Percentile | 888.2 | 1177 | 789.9 | 1090 | 888.7 | 1179 | 831.4 | 846.3 |
| Maximum | 2169 | 9277 | 2147 | 3169 | 4606 | 6115 | 1754 | 2435 |
| Mann-Whitney test | **** | | **** | | **** | | n.s. | |

**Area — 240 min**

| Cell line pair | A | | A | | B | | C | |
|---|---|---|---|---|---|---|---|---|
| Genotype | Abi3-WT | Abi3-KO | Abi3-WT | Abi3-KO | Abi3-WT | Abi3-KO | Abi3-WT | Abi3-KO |
| Number of values | 430 | 442 | 270 | 289 | 392 | 279 | 345 | 260 |
| Minimum | 94.81 | 89.15 | 43.96 | 199.4 | 92.73 | 21.93 | 22.51 | 144.7 |
| 25% Percentile | 410.0 | 520.4 | 438.3 | 606.4 | 461.7 | 721.8 | 440.3 | 509.1 |
| Median | 517.1 | 752.6 | 565.6 | 799.1 | 578.5 | 1021 | 565.0 | 684.7 |
| 75% Percentile | 673.0 | 1010 | 721.7 | 1044 | 749.5 | 1447 | 727.2 | 875.9 |
| Maximum | 1697 | 3905 | 2017 | 2596 | 3152 | 6402 | 3359 | 6415 |
| Mann-Whitney test | **** | | **** | | **** | | **** | |

**Solidity — 15 min**

| Cell line pair | A | | A | | B | | C | |
|---|---|---|---|---|---|---|---|---|
| Genotype | Abi3-WT | Abi3-KO | Abi3-WT | Abi3-KO | Abi3-WT | Abi3-KO | Abi3-WT | Abi3-KO |
| Number of values | 221 | 287 | 286 | 140 | 613 | 517 | 449 | 552 |
| Minimum | 0.6740 | 0.6370 | 0.5630 | 0.6710 | 0.6480 | 0.5160 | 0.5950 | 0.6610 |
| 25% Percentile | 0.8285 | 0.8380 | 0.8310 | 0.8903 | 0.8415 | 0.8660 | 0.7975 | 0.8530 |
| Median | 0.8620 | 0.8770 | 0.8810 | 0.9125 | 0.8840 | 0.9000 | 0.8540 | 0.8890 |
| 75% Percentile | 0.8920 | 0.9050 | 0.9245 | 0.9300 | 0.9160 | 0.9260 | 0.8980 | 0.9188 |
| Maximum | 0.9500 | 0.9510 | 0.9680 | 0.9690 | 0.9630 | 0.9650 | 0.9690 | 0.9590 |
| Mann-Whitney test | ** | | **** | | **** | | **** | |

**Solidity — 30 min**

| Cell line pair | A | | A | | B | | C | |
|---|---|---|---|---|---|---|---|---|
| Genotype | Abi3-WT | Abi3-KO | Abi3-WT | Abi3-KO | Abi3-WT | Abi3-KO | Abi3-WT | Abi3-KO |
| Number of values | 385 | 402 | 291 | 227 | 595 | 538 | 454 | 429 |
| Minimum | 0.2900 | 0.3290 | 0.3240 | 0.3850 | 0.3130 | 0.3930 | 0.3160 | 0.4250 |
| 25% Percentile | 0.5905 | 0.7170 | 0.5670 | 0.7290 | 0.6320 | 0.6958 | 0.6550 | 0.7030 |
| Median | 0.7070 | 0.8260 | 0.6740 | 0.8100 | 0.7320 | 0.7810 | 0.7640 | 0.7720 |
| 75% Percentile | 0.7950 | 0.8920 | 0.7880 | 0.8870 | 0.8100 | 0.8600 | 0.8313 | 0.8390 |
| Maximum | 0.9300 | 0.9650 | 0.9340 | 0.9600 | 0.9520 | 0.9630 | 0.9590 | 0.9560 |
| Mann-Whitney test | **** | | **** | | **** | | * | |

**Solidity — 120 min**

| Cell line pair | A | | A | | B | | C | |
|---|---|---|---|---|---|---|---|---|
| Genotype | Abi3-WT | Abi3-KO | Abi3-WT | Abi3-KO | Abi3-WT | Abi3-KO | Abi3-WT | Abi3-KO |
| Number of values | 432 | 422 | 272 | 261 | 392 | 392 | 324 | 359 |
| Minimum | 0.2270 | 0.2070 | 0.2230 | 0.2750 | 0.1900 | 0.2800 | 0.2060 | 0.2760 |
| 25% Percentile | 0.5840 | 0.5908 | 0.4763 | 0.5445 | 0.4790 | 0.5420 | 0.4768 | 0.5400 |
| Median | 0.6830 | 0.7555 | 0.5835 | 0.6570 | 0.6200 | 0.6595 | 0.5985 | 0.6650 |
| 75% Percentile | 0.7920 | 0.8663 | 0.7120 | 0.8100 | 0.7308 | 0.7768 | 0.7295 | 0.7870 |
| Maximum | 0.9660 | 0.9670 | 0.9430 | 0.9650 | 0.9360 | 0.9520 | 0.9500 | 0.9600 |
| Mann-Whitney test | **** | | **** | | **** | | **** | |

**Solidity — 240 min**

| Cell line pair | A | | A | | B | | C | |
|---|---|---|---|---|---|---|---|---|
| Genotype | Abi3-WT | Abi3-KO | Abi3-WT | Abi3-KO | Abi3-WT | Abi3-KO | Abi3-WT | Abi3-KO |
| Number of values | 430 | 442 | 270 | 289 | 392 | 279 | 345 | 260 |
| Minimum | 0.2680 | 0.1550 | 0.2510 | 0.2040 | 0.1970 | 0.2140 | 0.2010 | 0.2400 |
| 25% Percentile | 0.5875 | 0.5430 | 0.5295 | 0.4855 | 0.4348 | 0.5350 | 0.4275 | 0.5315 |
| Median | 0.6965 | 0.6925 | 0.6505 | 0.6410 | 0.5445 | 0.6570 | 0.5520 | 0.6655 |
| 75% Percentile | 0.7945 | 0.8150 | 0.7655 | 0.8020 | 0.6878 | 0.7740 | 0.6805 | 0.7880 |
| Maximum | 0.9410 | 0.9630 | 0.9470 | 0.9620 | 0.9510 | 0.9490 | 0.9440 | 0.9650 |
| Mann-Whitney test | n.s. | | n.s. | | **** | | **** | |

***Supplementary Table 1. Summary of the statistical analysis performed on Abi3-WT and -KO macrophages following the different repetitions of the spreading assay.*** *The spreading assay was repeated three times, using three different pairs of cell lines (Cell line pair A-C). For each pair, the table shows percentiles values of each line at all the time points (15-240 min) for both area and solidity values, as well as the number of cells evaluated in each case. The results of the Mann-Whitney test performed on each pair of cell lines are reported on the table by means of asterisks; *$p \leq 0.05$, ** $p \leq 0.01$, *** $p \leq 0.001$, ****$p \leq 0.0001$.*

| | Segment length | | Segment mean diameter | | Segment volume | | Total no. branching points | | Full branch level | | Sholl intersections | | Sholl A.U.C. | | Cell area | | Cell volume | |
|---|---|---|---|---|---|---|---|---|---|---|---|---|---|---|---|---|---|---|
| | t value | p value | t value | p value | t value | p value | t value | p value | t value | p value | t value | p value | t value | p value | t value | p value | t value | p value |
| **Sex** | 1.214 | 0.238 | 0.448 | 0.658 | 0.675 | 0.507 | -0.354 | 0.727 | 0.94 | 0.358 | -0.521 | 0.602 | 0.636 | 0.526 | 0.03 | 0.977 | -0.144 | 0.8865 |
| **Genotype** | 8.251 | 3.12e-08 | 5.991 | 6.01e-06 | 6.974 | 6.83e-07 | -23.727 | <2e-16 | -10.98 | 3.65e-10 | -23.383 | $<2e^{-16}$ | -25.171 | $<2e^{-16}$ | -12.53 | 3.29e-11 | -2.657 | 0.0147 |

***Supplementary Table 2. Summary of the t- and p-values calculated for the fixed effects included in the linear mixed-model used for the statistical analysis of the data generated from Imaris.*** *The results of the 3D tracing of single microglial cells were analysed using a linear mixed-model including genotype and sex as fixed effects and mouse ID as random effect; wherever appropriate, branch level or Sholl radius were included as an additional fixed effect (data not shown). The distributions of the outcome variables were visually inspected and, if skewed, log-transformed. The model fits were visually inspected with help of QQ-plots (data not shown). The reported t- and p-values represent the effect size and statistical significance, respectively, for the fixed effects. A positive t-value corresponds to a higher mean of the outcome variable in Abi3-KO mice as compared to Abi3-WT, whereas a negative sign indicates a smaller mean in Abi3-KO mice.*

| Radius | Sex | | Genotype | |
|---|---|---|---|---|
| | t value | p value | t value | p value |
| **3** | -0.968 | 0.34420 | -3.300 | 0.00341 |
| **6** | -0.055 | 0.957 | -6.423 | $2.29e^{-06}$ |
| **9** | -0.797 | 0.434 | -10.316 | $1.12e^{-09}$ |
| **12** | 0.779 | 0.445 | -11.474 | $1.66e^{-10}$ |
| **15** | 0.217 | 0.829 | -14.442 | $<2e^{-16}$ |
| **18** | 1.229 | 0.233 | -14.079 | $3.63e^{-12}$ |
| **21** | 2.257 | 0.0259 | -15.963 | $<2e^{-16}$ |
| **24** | -0.23 | 0.819 | -15.19 | $<2e^{-16}$ |
| **27** | -0.079 | 0.938 | -13.198 | $2.03e^{-11}$ |
| **30** | 0.661 | 0.517 | -10.707 | $2.77e^{-09}$ |
| **33** | -0.883 | 0.388 | -10.285 | $6.50e^{-10}$ |
| **36** | -2.300 | 0.0242 | -5.947 | $7.78e^{-08}$ |
| **39** | -1.063 | 0.29132 | -3.891 | 0.00023 |
| **42** | -1.167 | 0.2486 | -2.417 | 0.0194 |
| **45** | -0.560 | 0.5798 | -2.024 | 0.0523 |
| **48** | 0.222 | 0.82669 | -1.159 | 0.26156 |
| **51** | -0.115 | 0.911047 | -2.673 | 0.026531 |
| **54** | 0.544 | 0.6061 | -1.269 | 0.2497 |

***Supplementary Table 3. Summary of the t- and p-values obtained for the Genotype and Sex effect for each radius of the Sholl analysis.*** *The Sholl radius was included in the model as a fixed effect (data not shown) to account for the variability in branch number related to the distance from the centre.*

***Supplementary Video1. Representative time stack of microglia movement in awake ABI3-WT mice as observed through two-photon microscopy.*** *Each of the 30 time points comprising the time stack represents the orthogonal projection of a 50µm-thick z-stack. Z-stacks were acquired every 90 seconds over a 45-minute-long imaging session using a 7x zoom in order to better visualise processes*.

***Supplementary Video2. Representative time stack of microglia movement in awake ABI3-KO mice as observed through two-photon microscopy.*** *Each of the 30 time points comprising the time stack represents the orthogonal projection of a 50µm-thick z-stack. Z-stacks were acquired every 90 seconds over a 45-minute-long imaging session using a 7x zoom in order to better visualise processes.*
